# Apparent increase in lip size is linked to tactile discrimination improvement

**DOI:** 10.1101/855296

**Authors:** Elisabetta Ambron, H. Branch Coslett

**Author notes:** Corresponding author: Dr. Elisabetta Ambron, Dept. of Psychological & Brain Sciences, University of Delaware, 77 E. Delaware Avenue, Newark, DE 19711.

## Abstract

Magnified vision of one’s body part has been shown to improve tactile acuity. We used an anesthetic cream (AC) to explore this effect. In Experiment 1, application of AC caused an increase in perceived lip size. As perceived lip size increased, tactile discrimination (as assessed by two-point discrimination) improved. In Experiment 2, we replicated these results in a larger sample and showed that these effects are not observed without AC. Tactile discrimination was better and improved as a function of lip size with AC, while performance remained consistent without AC. In Experiment 3, we showed that the increase in perceived lip size occurred only with the application of AC, but not with moisturizing cream. The application of either cream induced an improvement in two-point discrimination, but this improvement was modulated by perceived lip size only for AC. Magnification effects are mediated by malleable, experience-dependent representations of the human body.

## Introduction

Multiple sources of information, including sensory and vestibular inputs as well as feedback from action, are integrated to generate a representation of the body that specifies the position and size of body parts^1,2,3^. Evidence for the dynamic multisensory nature of this representation comes from demonstrations that the perceived size of a body part can be increased by modifying visual^3–7^ or somatosensory inputs^2, 8–9^

Several investigators have demonstrated that changes in the perceived size of a body part may have consequences for sensory^4^ or motor function^6^. For instance, Kennett and colleagues^4^ showed that performance on a two-point discrimination task involving the forearm improved when looking at the forearm through magnifying lenses as compared to normal vision (magnification effect). Additionally, we have demonstrated that magnifying lenses increase motor cortex excitability in normal subjects^10^ and improve motor function in subjects with stroke^11^.

Although previous work has explored the effects of visual perturbations of body size on sensory and motor function, we are unaware of investigations exploring the effects of magnification of a body part achieved by non-visual manipulations. If the effect of magnification is mediated by a higher order body representation that integrates multiple sensory modalities^3^, one would predict that improved tactile discrimination would be observed when body parts are perceived as larger.

To test this hypothesis, we exploited the report by Gandevia and Phegan^8^ that application of an anesthetic cream (AC) to the lips was associated with an increase in perceived lip size without complete anesthetization of the lips. We replicated these findings in pilot testing (n=4) using a different anesthetic (benzocaine), finding that participants experienced an increase in lip size while still being able to perform two-point discrimination. Given this finding, we used AC to examine our primary question of interest. We presented individuals with a two-point discrimination task on the lips in three conditions: before application of the AC, shortly after the AC was applied (when the lips should be perceived as enlarged), and after the AC was removed and perceived lip size returned close to normal. Full anesthetization of the lips was not induced, but the application of the cream was enough to induce the sensation of an increase in lip size in most participants. If local anesthetics induce the perception that the treated body part is larger, and this representation of the body underlies performance on a two-point discrimination task, one would expect that subjects who feel larger lips would perform better at a tactile task such as two-point discrimination. We predicted that the improvement in tactile discrimination should be observed in the more ambiguous conditions, such as when participants were presented with 2 stimuli at a short distance (1 or 2 mm). On the contrary, we expected the identification of 1-point stimulation (0mm distance) and of 2 points with a larger separation in space (3 mm) would be easier and less likely to show an improvement. We tested this prediction in three experiments.

In Experiment 1, consistent with our hypothesis, there was a significant improvement on the tactile task with AC and this improvement was related to the magnitude of the perceived increase in lip size. These results were replicated in a larger sample in Experiment 2. Thus, despite the fact that participants’ lips were partially “numb”, two-point discrimination was improved with AC; these data suggest that AC has differential effects on nociception and other aspects of somatosensory processing and that changes in the latter underline the perception of lip size and the enhancement of two-point discrimination. In Experiment 2 we found a significant relationship between the increase in perceived lip size and tactile discrimination only when AC was applied, and not in the control condition without the application of the cream. Based on previous work regarding the mechanism of action of local anesthetics^12^, we propose that the perception of lip enlargement observed with AC may result in an increase of the cortical sensory representation of the lip.

One possibility is that the effects observed in Experiments 1 and 2 are a non-specific effect of the application of a cream. We note that Lévêque et al.^13^ reported an improvement in two-point discrimination with skin hydration in older subjects, whereas Guest et al.^14^ did not find an effect of moisturizing cream on a grating orientation task with young subjects. To address the possible non-specific effects of AC, in Experiment 3 we examined whether the improvement in two-point discrimination observed in Experiments 1 and 2 was linked to changes in perceived lip size or could be induced by a moisturizing cream (EucerinTM, hereafter MC). Based on the findings above, we had no strong prediction regarding the effect of MC on overall performance but, crucially, expected to observe a relationship between perceived changes in lip size and two-point discrimination for the AC but not for MC. If AC increases the perceived size of the body, and this change in body size causes improved tactile performance, one would expect a significant positive relationship between tactile performance and subjective enlargement of the lips. In contrast, in the MC condition we predicted no significant changes in perceived lip size and, therefore, no significant effect of changes in perceived lip size on two-point discrimination performance. Our predictions were confirmed.

## Results Experiment 1

First, we tested whether AC was effective in inducing a change in the perceived lip size. As shown in Table 1, participants rated the size of their lips bigger after the application of AC (*t*=5.2, *p*<0.001). This sensation of the increase in lip size decreased once the cream was removed (*t*=5.2, *p*<0.001), so that participants’ ratings of the lip size in the POST were similar to PRE (*t*=0.69, *p*=0.49) condition.

**Table 1.** Means (SEM) for lip size, numbness and tactile detection test across conditions for the three experiments Same symbols indicate significant differences (*p*<0.05) between conditions.

|  |  | ANESTHETIC |  |  | CONTROL CONDITION |  |  |  |
| --- | --- | --- | --- | --- | --- | --- | --- | --- |
|  |  | PRE | EXP | POST | PRE | EXP | POST | Omnibus |
| **EXPERIMENT 1** | *Lip size –lips only* | 4.2(0.4)* | 5.7 (0.4)*# | 4(0.4)# | NA | NA | NA | Time $F(2,36) = 17.14, p<0.001, \eta^2=0.49$ |
| | *Lip size – whole face* | 2.9 (0.2)* | 4.4 (0.3)*# | 3.05 (0.2)# | NA | NA | NA | Time $F(2,36) = 24.3, p<0.001, \eta^2=0.57$ |
| **EXPERIMENT 2** | *Lip size –lips only* | 4.4(0.3)*§ | 5.8(0.3)*# | 4.9(0.3)#§ | 4.3(0.3) | 4.3(0.3) | 4.4(0.3) | Time $F(2,78) = 27.6, p<0.001, \eta^2=0.41$<br>Cream $F(1,39) = 24.3, p<0.001, \eta^2=0.38$<br>Cream x Time $F(2,78) = 30.8, p<0.001, \eta^2=0.44$ |
| | *Lip size – whole face* | 2.7 (0.2)*§ | 3.9 (0.2)*# | 3.4 (0.2)#§ | 2.9 (0.2) | 2.8 (0.2) | 2.9 (0.2) | Time $F(2,78) = 21, p<0.001, \eta^2=0.36$<br>Cream $F(1,39) = 30, p<0.001, \eta^2=0.44$<br>Cream x Time $F(2,78) = 31, p<0.001, \eta^2=0.45$ |
| | *Numbness* | 1.6(0.3)*§ | 6.9(0.3)*# | 5.2(0.4) #§ | 1.2(0.3)^ | 1.8(0.3)^ | 1.7(0.0) | Time $F(2,78) = 81, p<0.001, \eta^2=0.72$<br>Cream $F(1,39) = 102, p<0.001, \eta^2=0.67$<br>Cream x Time $F(2,78) = 51.4, p<0.001, \eta^2=0.56$ |
| | *Tactile Detection* | 0.72 mN (0.23) | 0.66(0.19) | 0.61(0.13) | 0.47 mN (0.14) | 0.61 mN (0.19) | 0.73(0.23) | Time $F(2,78) = 0.10, p=0.75, \eta^2=0.003$<br>Cream $F(1,39) = 0.16, p=0.85, \eta^2=0.004$<br>Cream x Time $F(2,78) = 0.89, p=0.41, \eta^2=0.023$ |
| **EXPERIMENT 3** | *Lip size– lips only* | 3.6 (0.4)*§ | 5.5 (0.4)*# | 4.6 (0.3)#§ | 3.7 (0.4) | 4 (0.3) | 3.7 (0.4) | Time $F(2,26) = 11, p<0.001, \eta^2=0.46$<br>Cream $F(1,13) = 5.9, p=0.03, \eta^2=0.31$<br>Cream x Time $F(2,26) = 8.9, p<0.001, \eta^2=0.72$ |
| | *Lip size– whole face* | 2.7 (0.2)*§ | 4.1 (0.2)*# | 2.4 (0.3)#§ | 3 (0.3) | 3.2 (0.2) | 2.9 (0.3) | Time $F(2,26) = 11, p<0.001, \eta^2=0.46$<br>Cream $F(1,13) = 4.6, p=0.05, \eta^2=0.26$<br>Cream x Time $F(2,26) = 11.4, p<0.001, \eta^2=0.047$ |
| | *Numbness* | 1.4 (0.7) *§ | 6 (0.6)*# | 4.6 (0.4)#§ | 1.1 (0.4) | 1.5 (0.4) | 1.4 (0.4) | Time $F(2,26) = 19.8, p<0.001, \eta^2=0.60$<br>Cream $F(1,13) = 21, p=0.001, \eta^2=0.61$<br>Cream x Time $F(2,26) = 11, p<0.001, \eta^2=0.46$ |
| | *Tactile Detection* | 0.44 mN (0.10) | 1.8 mN (0.9) | 0.57 mN (0.9) | 0.46 mN (0.10) | 0.19 mN (0.10) | 0.26 mN (0.14) | Time $F(2,26) = 0.39, p=0.63, \eta^2=0.03$<br>Cream $F(1,13) = 3.2, p=0.09, \eta^2=0.20$<br>Cream x Time $F(2,26) = 1.0, p=0.35, \eta^2=0.08$ |

Second, we conducted a logistic mixed effect model (LMM) analysis including time point (PRE, AC, and POST) and distance (0, 1, 2 and 3 mm) as fixed categorical factors. Distance was defined as a categorical factor in order to investigate whether the improvement occurred specifically at 1 or 2 mm.

The final model included both factors (logLik= −2337, χ^2^ (8)=58.1, *p*<0.001). As shown in Table 1 SI, the main effects of the distance and the interaction time by distance were significant. In line with our predictions, participants identified two point stimulation more frequently in the AC compared to the PRE when the distance was 1 (*z*=-3.3, *p*<0.001) and 2 mm (*z*=-3.7, *p*<0.001), but also at 3 mm (*z*=-3.4, *p*<0.001) (see Figure 1). Performance was worse at 0mm distance in the AC than PRE (*z*=2.4, *p*=0.01). Similar performance was observed in the AC and POST across all distances, except for 1mm (*z*=-2.4, *p*=0.01). This evidence suggests that a learning effect might have played an additional role in our results.

**Figure 1:**
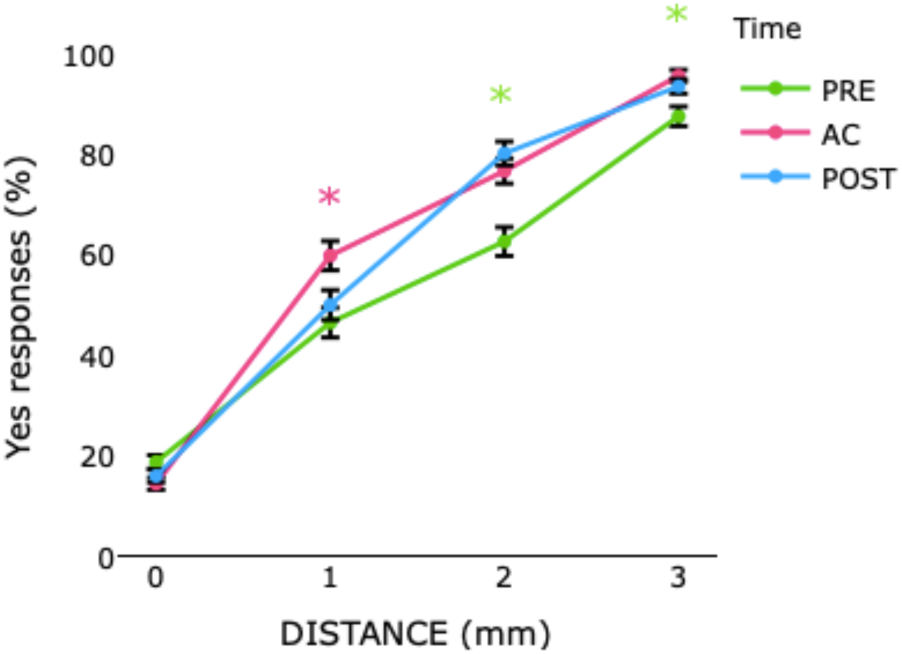
Mean (markers) and SE of the means (error bars) of yes responses in two points discrimination in PRE, AC and POST conditions for each distance (0,1,2,3 mm) in Experiment 1. The start sign indicates significant (*p*<0.05) effect of the increase in the comparison across conditions. The color of the star indicates the which condition differed from the others

These results provide clear evidence that administration of AC paradoxically improves performance on a two-point discrimination task. Is there a direct relationship between the amount of perceived lip enlargement and the improvement in two-point discrimination? To further investigate this, we determined the change in perceived lip size in the AC and POST conditions compared to the PRE condition. This was scored as the difference between the rating of perceived lip size in the PRE and the AC or POST conditions. For example, if a subject rated their lip size as corresponding to the 3^rd^ face on the eight-point scale in the PRE condition and the 5^th^ face for the CREAM condition, the difference score was +2.

Next we ran two LMMs for number of *yes* responses on two points stimulation for the AC and the POST conditions, with change in perceived lip size as continuous and distance (0, 1, 2 and 3 mm) as categorical fixed factors. To control for the possible influence of basic tactile abilities, we inserted the *yes* responses at two points stimulation in the PRE condition as a fixed factor in the model, but it did not contribute significantly to the model and was removed. For the AC condition, the final model included both factors: distance and change in perceived lip size – whole face measure (logLik= −668, χ^2^ (4)=9.2, *p*=0.05). The main effects of both distance and lip size were significant (see Table 1 SI for the omnibus of the effects). Although the interaction did not reach significance, we tested our prediction that the increase in the perceived lip size induced changes in the *yes* responses in particular when there was a separation between the two points of 1 and/or 2 mm. In line with our hypothesis we observed a significant increase of *yes* responses as function of the increase in the perceived lip size for 1mm (*z*=3.1, *p*=0.001) and 2mm (*z*=2.7, *p*=0.02) distances (see Figure 2 left panel). When the cream was removed and the sensation of increased lip size disappeared in the POST condition, distance was the only significant predictor of *yes* responses (logLik= −770, χ^2^ (3)=871, *p*<0.001) (see Table 1 SI and Figure 2 right panel). This evidence suggests that although a learning effect might have been present, it could only partially account for the results observed in the AC condition.

**Figure 2:**
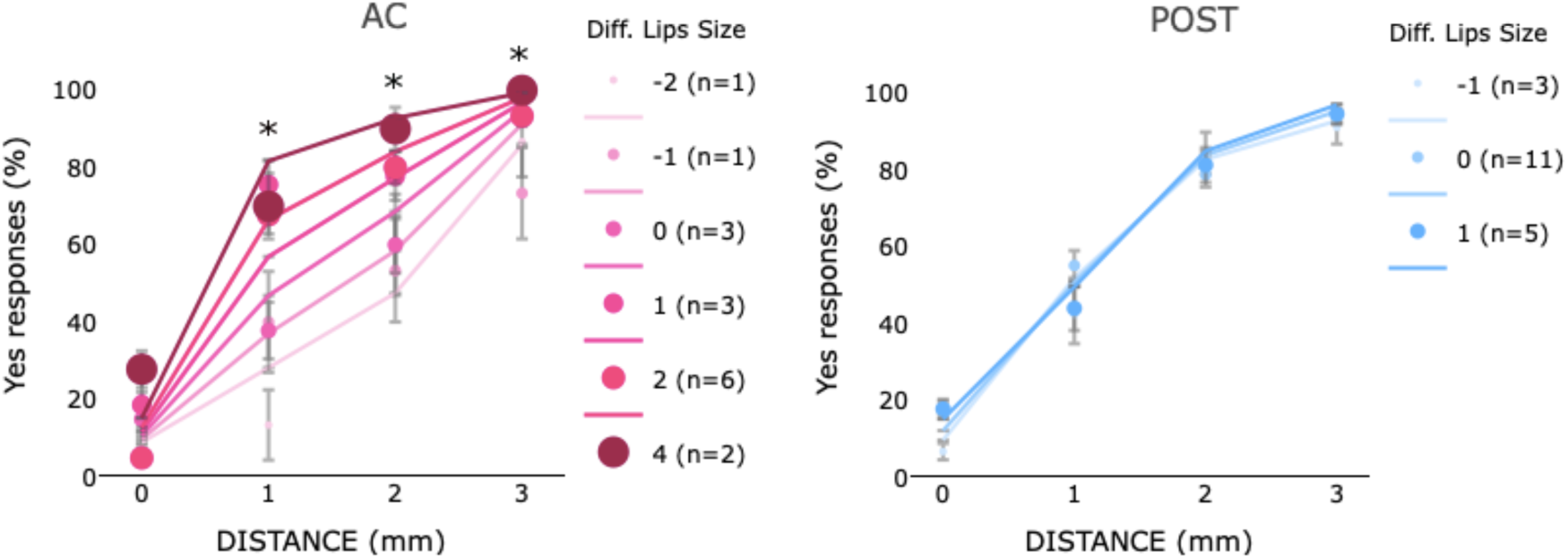
Mean (markers) and SE of the means (error bars) of yes responses in two point discrimination in AC (left panel) and POST (right panel) conditions as a function of distance (0,1,2,3 mm) between 2 stimulation sites and the change in the perceived lip size in Experiment 1. The different size/coloring of the markers described the performance of individuals based on change in the perceived lip size in that condition with respect to the PRE. The legend describes the amount of perceived change in the lip size and the number of individuals who perceived that change. Negative numbers indicate a perceived decrease and positive numbers indicate an increase in the perceived lip size compared to PRE. For example, ‘-1 (n=1)’ indicates that one participant perceived a decrease in lip size of 1 unit, while ‘1 (n=5)’ indicates that five participants experienced an increase in lip size of 1 unit. In every plot, colored lines represent the estimated effects in the LMM analyses.

## Results Experiment 2

This experiment was carried out to replicate the results of Experiment 1 with a larger sample and to compare the application of AC with a control condition in which there was no application of the cream. Additionally, we sought to formally evaluate the degree of anesthesia produced by the AC by obtaining ratings of numbness using a rating scale and objective measures of tactile detection (Semmes-Weinstein monofilament, Stoelting).

First, we examined whether or not AC increased perceived lip size and changed the sensation of numbness without substantially altering tactile detection and whether these effects were observed without the application of AC. As shown in Table 1, we found significant main effects and interactions for the lip size and numbness ratings, but not for tactile detection as assessed by Semmes-Weinstein monofilament. Indeed, participants rated the size of the lips bigger and the level of numbness greater in the EXP than PRE condition for the AC condition. An increase of numbness was also noted in the EXP than PRE condition for the no AC condition. To follow up these results, we computed the difference between EXPERIMENT and PRE conditions for both lip size measures and numbness. The differences were larger in the AC than no AC condition for both lip size measures (lips only score *t*(39)=6.9, p<0.001; whole face score *t*(39)=6.7, p<0.001) and numbness, *t*(39)=9.9, *p*<0.001.

Second, we examined whether AC induced a change in two-point discrimination compared to the no AC condition. We ran a LMM analysis with the categorical factors condition (AC or no AC), time point (PRE, EXPERIMENT, and POST) and distance (0, 1, 2 and 3 mm); all were included in the final model (logLik= −10550, χ^2^ (12)=49.3, *p*<0.001). As shown in Table 1 SI, the 3-way interaction was marginally significant: in line with our predictions, participants showed a better performance with AC than no AC with 1 (*z*= −3.5, *p*<0.001) and 2mm (*z*= −3.7, *p*<0.001) distance, while similar performance between conditions was observed at 0 and 3 mm (see Figure 3).

**Figure 3:**
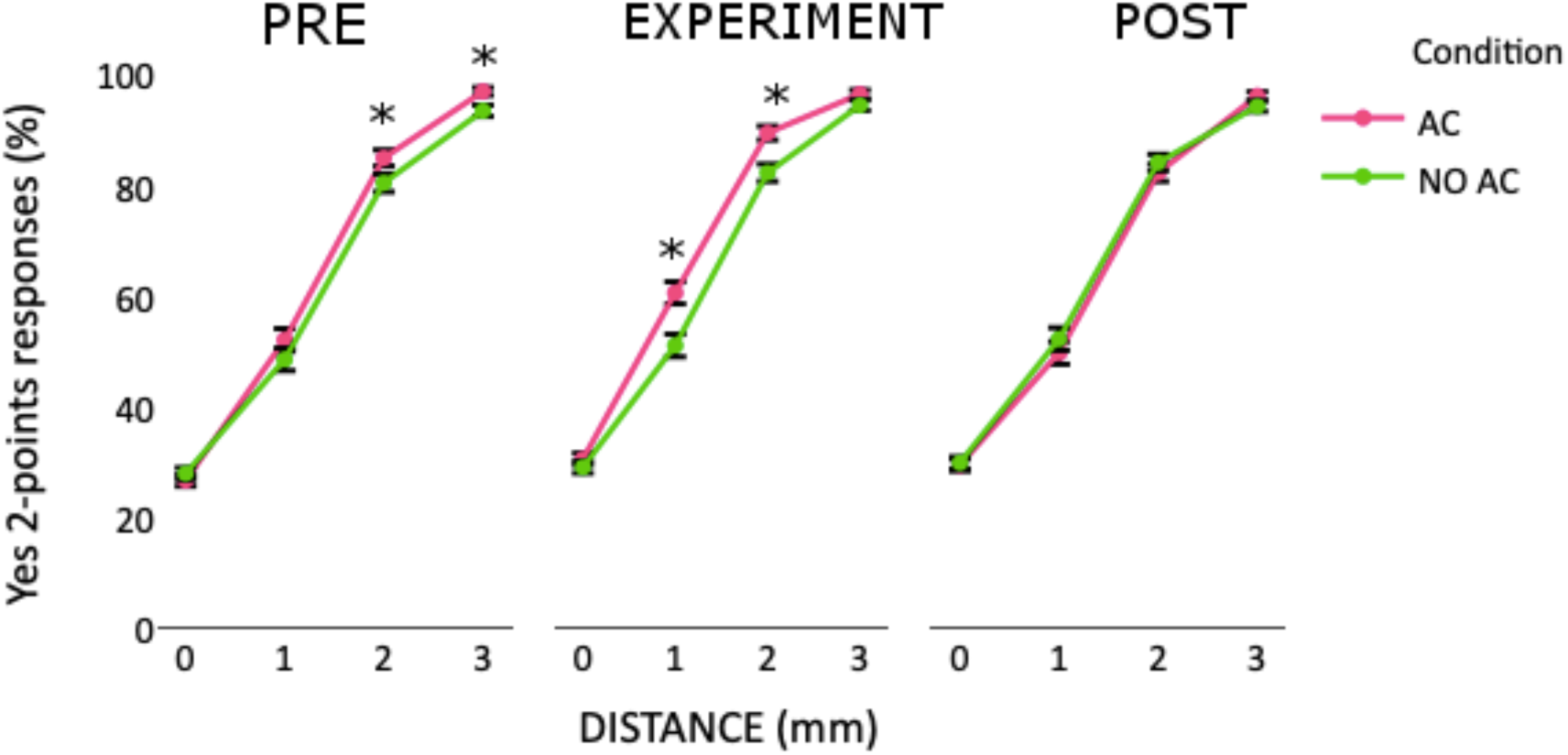
Mean (markers) and SE of the means (error bars) of yes responses in two point discrimination in PRE, EXPERIMENT and POST conditions for AC and no AC conditions at each distance (0,1,2,3 mm) in Experiment 2.

Furthermore, we expected to replicate the results of Experiment 1 showing that *yes* responses increase with the AC application. We also predicted that these effects would not be observed in the no AC condition. In line with our predictions, performance in the EXPERIMENT condition was better than PRE condition at all distances (*z*>2.3, *p*<0.05) except for 3 mm (*z*=0.49, *p*=0.61), while performance remained consistent for the no AC condition.

To test if the performance at baseline could account for the observed difference in the EXPERIMENT condition, we carried out an additional LMM analysis on this condition inserting the *yes* responses at baseline as a fixed factor. This factor did not improve the model fit with respect to the model with distance and condition as fixed factor (logLik= −3534, χ^2^ (8)=10.8, *p*=0.20) and its inclusion did not change our results, suggesting that the variability in the baseline performance across condition could not account for our results.

Third, we investigated the effect of the change in the perceived lip size on tactile discrimination by computing the difference in perceived lip size in the EXPERIMENT condition compared to the PRE condition. We then examined how the change in perceived lip size, along with distance, affected *yes* responses in both no AC and AC condition. While for no AC the final model included only the fixed factor distance (logLik= −1768, χ^2^ (3)=1273, *p*=0.001), for the AC condition both change in lip size (lips only) and distance contributed to the final model (logLik= −1733, χ^2^ (6)=12.3, *p*=0.055) (see Figure 4). The main effects of distance and size were significant (see Table 1 S!I), suggesting that the *yes* responses increased with the distance, as well as with the perceived increase in lip size. In order to investigate our predictions, we tested whether the increase in lip size had a specific effect at 1 and/or 2mm. *Yes* responses increased with the increase in the lip size at distance 2mm (*z*=1.9, *p*=0.054) and also at 0 mm (*z*=2.3, *p*=0.01), but not at 1mm or 3 (*z*<0.9, *p*>0.39).

**Figure 4:**
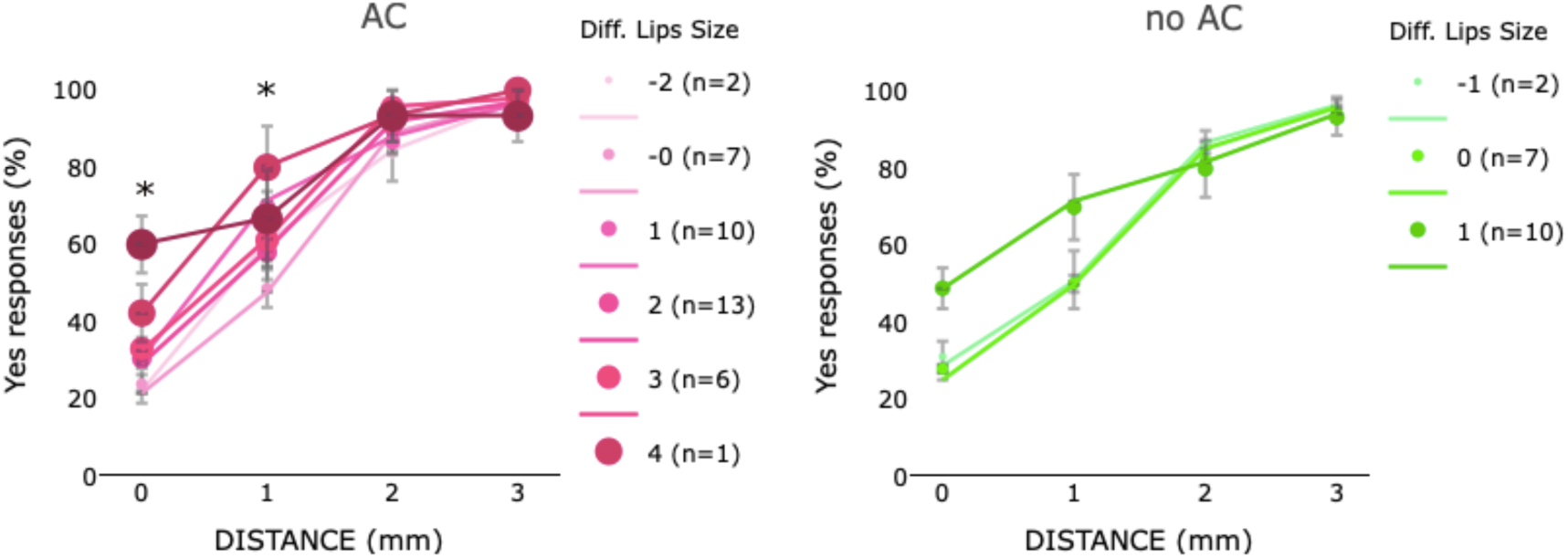
Mean (markers) and SE of the means (error bars) of yes responses in two points discrimination in AC (left panel) and no AC (right panel) conditions as a function of the distance (0,1,2,3 mm) between 2 stimulation sites and the change in the perceived lip size (lips only rating) in Experiment 2. The different size/coloring of the markers described the performance of individuals based on change in the perceived lip size in that condition with respect to the PRE. The legend describes the amount of perceived change in the lip size and the number of individuals who perceived that change. Negative numbers indicate a perceived decrease and positive numbers indicate an increase in the perceived lip size compared to PRE. In every plot, colored lines represent the estimated effects in the LMM analyses.

## Results Experiment 3

Experiment 3 was performed to control for the possibility that changes in perceived lip size are a non-specific effect of AC that can be observed when applying any cream to the lips.

First, we explored whether or not AC and MC induced changes in perceived lip size, numbness and tactile detection using multiple repeated measures ANOVAs with time points (PRE, EXPERIMENT, POST) and condition (MC, AC) as factors. As shown in Table 1 there were no significant main effects or interactions for tactile detection, demonstrating that the anesthetization did not significantly influence tactile detection. However, significant main effects and interactions were observed for the perceived lip size and numbness. Post hoc t-tests were carried comparing difference (AC-PRE) of size and numbness between AC and MC conditions. For the lip size measure, the difference was larger in AC than MOISR condition (lips only score *t*(13)=5.0, p<0.001; whole face score *t*(13)=4.6, *p*<0.001). Similarly, the difference in numbness was higher in the AC than MC, *t*(39)=3.3, *p*=0.005).

Second, we investigated if two-point discrimination performance differed with the application of AC or MC. We ran a LMM analysis with cream type (AC or MC), time point (PRE, EXPERIMENT, and POST) and distance (0, 1, 2 and 3 mm) as fixed categorical factors and all were included in the final model (logLik= −3487, χ^2^ (12)=49, *p*<0.001). As shown in Table 1 SI, the three-way was significant. In the EXPERIMENT condition participants were more accurate in the MC than AC at 1,2,3 mm distances (*z*>2.8, *<u>p</u>*<0.01) (see SI Figure 3).

Also, we were expected to replicate the results of Experiment 1 and 2, consisting in an increase in *yes* responses with the application of AC compared to the PRE condition at 1 and/or 2 mm. This prediction was met as we observed an improvement at 1mm (*z*=2.05, *p*<0.03). The application of MC also induced an improvement at 1,2,3 mm (*z*>2.8, *p*<0.01 in all comparison).

This evidence could suggest that the application of both cream induced an improvement and that similar principle could account for both cream application. To test this hypothesis, we carried out a LMM analysis for each cream (AC and MC), with changes in lip sizes coded as continue and distance (0, 1, 2 and 3) coded as categorical factor on *yes* responses. As the difference between AC and MC was primarily observed in the EXPERIMENT condition, we focused on this condition. Changed in lip size would predict *yes* responses in the EXPERIMENT condition for both AC and MC if the improvement observed with the application of both creams was based on similar mechanism. On the contrary, we expected changed in lip size to account for the *yes* response only for AC, if the increase in the perceived lip size was a specific mechanism of AC. In line with this last prediction, LMM analyses showed a significant contribution of the changes in lip size for the AC, but not for MC. Indeed, the model including both distance and lip size (whole face) better predicted *yes* responses than the model with only distance as fixed factor for the AC (logLik= −3487, χ^2^ (12)=49, *p*<0.001), but not for MC (logLik= −566, χ^2^ (4)=21, *p*<0.001). Although performance was better with MC than AC at 1 and 2 mm distances, the increase in perceived changes in lip size accounted for *yes* responses only for AC and not for the MC (see Figure 5). Indeed, *yes* responses increased as a function of perceived increase in lip size only for AC, at 2 mm (*z*=3, *p*=0.02) and 3 mm (*z*=2, *p*=0.01) distances (see left panel Figure 5).

**Figure 5:**
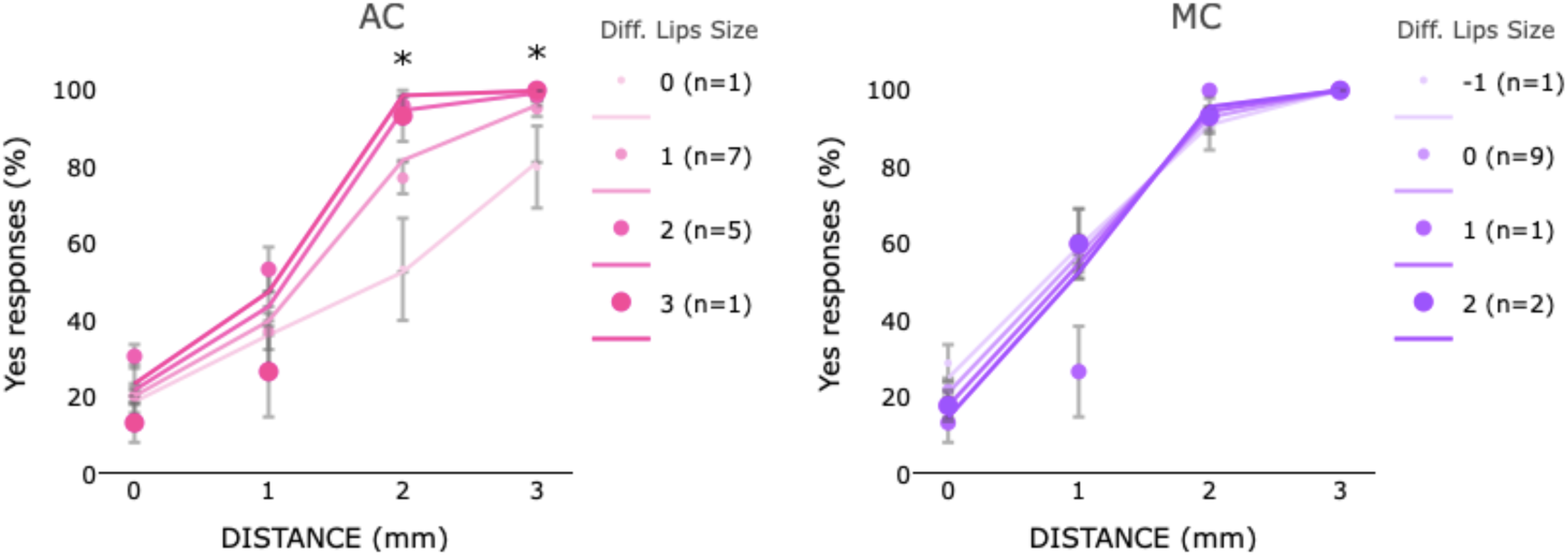
Mean (markers) and SE of the means (error bars) of yes responses in two points discrimination in AC (left panel) and MC (right panel) conditions as function of the distance (0,1,2,3 mm) between 2 points and the change in the perceived lip size (whole face rating) in Experiment 3. The different size/coloring of the markers described the performance of individuals based on change in the perceived lip size in that condition with respect to the PRE. The legend describes the amount of perceived change in the lip size and the number of individuals who perceived that change. Negative numbers indicate a perceived decrease and positive numbers indicate an increase in the perceived lip size compared to PRE. In every plot, colored lines represent the estimated effects in the LMM analyses.

## Discussion

Anesthetic cream applied to participants’ lips significantly improved two-point discrimination. This effect was observed in Experiment 1 and replicated in Experiments 2 and 3. In particular in Experiment 2 we replicated the results of Experiment 1 in a larger sample of participants (n=40) and demonstrated that the improvement observed in Experiment 1 was not simply an effect of learning, but was a result of the anesthetic. In Experiments 1 and 2, the magnitude of the effect was related to the participants’ judgment of the magnitude of increase in lip size. In Experiment 3, we also demonstrated that whereas both AC and MC improved two-point discrimination, only AC was associated with the perception of enlargement of the lips. To account for the present results, it is important to understand: (i) how AC and MC act at the peripheral level; (ii) how this mechanism might affect perceived body size; (iii) how these two factors interact to determine participants performance.

Benzocaine, a sodium channel blocker, primarily affects small, unmyelineted C-fibers responsible for pain sensation rather than large myelinated fibers involved in touch and proprioception^15^. As C-fibers typically inhibit the activity of mechanoreceptor inputs, reducing the activity of C-fibers is thought to induce an expansion of the receptive fields of mechanoreceptors^12, 13^. We suggest that the increase in receptive field size leads to improvement in two-point discrimination because it increases the precision with which the stimulated location can be marked on the skin surface. The putative mechanism underlying this effect invokes the principle of coarse coding^16^ and is illustrated in Figure 6. In the baseline (pre-cream) condition, the site stimulated falls within the receptive field of a single neuron (e.g. P2 in R4/N4); as stimulus site is defined only with respect to a single neuron, the site is indistinguishable from that of any other point in the area subserved by the neuron. As illustrated in Figure 6, with an increase in receptive field sizes, the same point could fall within the receptive fields of multiple neurons (e.g. P2 in R3/N3, R4/N4 and R5/N5), all of which respond to the stimulus. As the region of the skin subserved by the intersection of multiple mechanoreceptors is likely to be smaller than the area subserved by a single neuron in the absence of anesthetic, the site of the touch is marked with greater precision with AC, thereby enhancing the ability to distinguish between one and two points. We speculate that this account might also explain a seemingly paradoxical finding in the AC condition – that is, poorer performance on one-point trials. After application of AC, each stimulus would fall inside more receptive fields, inducing a response from a larger number of neurons than in the normal condition. An increase in the number of neurons responding to a single touch may, on one-point stimulation trials, lead to a bias to perceive that two sites were stimulated.

**Figure 6:**
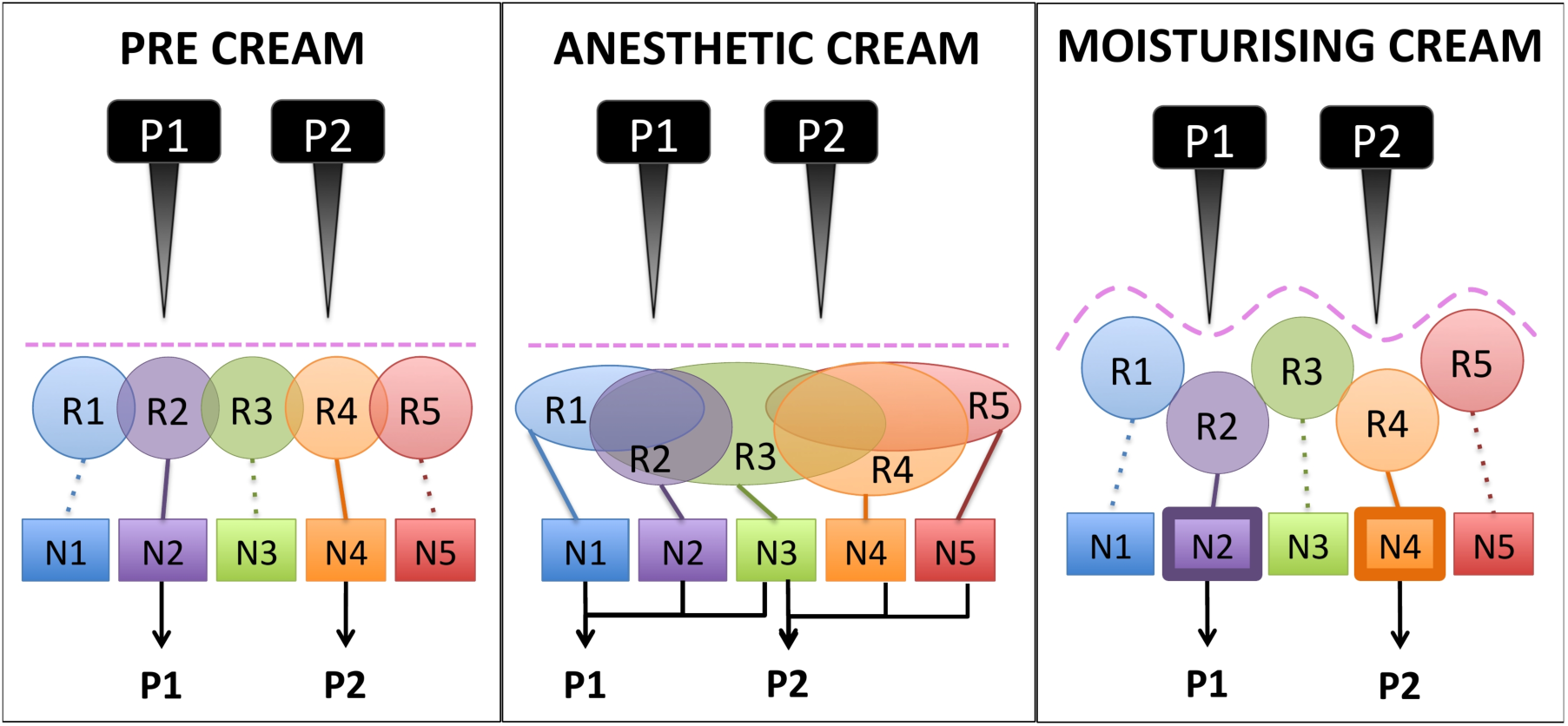
Schematic description of the possible changes occurring in the lips’ receptive fields in the PRE cream condition, with the anesthetic and moisturizing cream. R: receptive field of neuron in S1; N: neuron. P: points of stimulation 1 and 2.

We note that the account presented in Figure 6 is likely to be incomplete. Performance on a two-point discrimination task is likely to depend not only on the capacity to precisely discriminate between stimuli but also on lateral inhibition^17^. We are unaware of data regarding the effects of local anesthetics such as benzocaine on lateral inhibition.

Consistent with previous work demonstrating an improvement in two-point discrimination with skin hydration in elderly^13^, we observed improvement in two-point discrimination with MC in Experiment 3. This improvement, which was not associated with a perception of enlargement of the lips, is likely to be attributable to an entirely different mechanism from that observed with AC. As suggested by others^13^, we propose that MC increases skin volume and surface area, thereby increasing the physical distance between the skin’s mechanoreceptors (see Figure 6).

How does the putative increase in peripheral receptive field size lead to a change in the perceived size of the lips? We suggest that this is a reflection of the dynamic plasticity in the cortical representation of peripheral sensory inputs. Calford & Tweedale^12^ reported that local anesthetics induce a rapid increase in the receptive fields for neurons in SI corresponding to the anesthetized and immediately surrounding body surface. If the anesthetization does not completely silence the representation of that body part, neurons of both the anesthetized and surrounding areas will discharge, creating an increase in background noise that could induce the sensation of increased lip size^8^. We speculate that an increase in the size of the receptive fields of S1 neurons results in a perception that the lips are increased in size.

The present data may appear surprising given previous reports of a decrease in tactile discrimination^19^ and a reduction of skin sensation^28^ with local anesthesia. How can we reconcile this apparent contradiction? First, some previous studies used an anesthetic cream (ELMA), which has been shown to be more effective in reducing tactile sensitivity compared to the AC creams that we employed^29^. Second, we tested tactile discrimination a few minutes (∼ 2 minutes) after the cream application, while in previous work testing occurred approximately 30 minutes after application of the anesthetizing cream. It has been proposed that the decrease in tactile sensitivity may be the secondary effect of nociceptor blockage^29^ rather than a direct action of the local anesthetic on mechanoreceptors and cutaneous fibers. If so, it is possible that in our experiment, the type of cream used and/or time between the cream application and task performance was not sufficient for the anesthetic to be fully effective^18, 19^. Indeed, the results of Experiment 2 are consistent with this interpretation; we did not observe a significant reduction of tactile sensitivity as measured with the tactile detection task.

At a cortical level, deafferentation of a body part is thought to unmask cortical connections, increase the cortical excitability of the areas close to the anesthetized body part and induce cortical reorganization^20–21^. Interestingly, we have recently shown similar patterns of cortical changes with the increase in the perceived size of a visualized body part. We recorded motor evoked potentials while participants were looking at their hand with normal, unaltered vision or with a magnifying lens; we observed an increase in cortical excitability as measured by transcranial magnetic stimulation and of the size of the representation of the hand with magnification of the visual image of the hand^10^. We speculate that the use of the anesthetic cream may not only induce a comparable sensation of the increase in body part size, but also a cortical reorganization similar to that reported for the visual magnification of a body part. This hypothesis is also corroborated by evidence of an association between improvement in motor or tactile tasks and changes in cortical excitability, reported in studies that enhanced participants’ focus of attention on a specific body part^22^ or participants’ expectations regarding the effect of the experimental manipulation on their performances^23, 24^.

The visual increase of the size of a body part provides a beneficial effect on motor and somatosensory functions. Indeed, previous work has demonstrated that accuracy in two-point-discrimination tasks measured on the arm increases when looking at this body part with magnifying lenses^4^. We report for the first time that magnification of a body part achieved through manipulation of somatosensory input may improve two-point discrimination.

Although Calford & Tweedale^12^ reported an increase in receptive field size in S1 after lidocaine injection, we speculate that the effect of AC on the perceived size of the lips may be mediated by multimodal areas of the parietal cortex^5, 7, 25^. We previously argued for multiple discrete representations of the body^3^. Our account postulates that primary somatosensory representations coded in S1 articulate (bidirectionally) with the *body form representation* that codes the size and shape of the individual’s body. We argued that the *body form representation* is supported by multimodal parietal regions such as the anterior intraparietal sulcus. This area facilitates not only cross-modal effects between vision and touch^25^, but is also an important hub for multisensory integration^26^; changes in activity within this region may account for individual differences in multisensory integration^27^. Specifically, as the cortex of the anterior intraparietal sulcus mediates the beneficial effect of vision of the limb in tactile discrimination^25^, we speculate that this area might have an important role also in mediating the magnification effect across modalities.

A beneficial effect of local anesthesia on tactile detection and discrimination has also been noted in previous studies with normal adults^30^ and patients with nerve injuries^31^. However, these studies differ from our work, in that the cream was not applied on the same site of the tactile stimulation, but on body areas (e.g. the hand) close to the anesthetized body region (e.g. forearm). The idea behind this manipulation was that the application of the cream on the forearm would produce deafferentation and a reduction of the cortical representation of this body part with a compensatory increase of the cortical representation of adjacent body areas that could cause improvement in tactile discrimination tested in this region^31, 32^. The improvement observed in the present study is the result of the direct manipulation of perceived body size at the site of stimulation and may be mediated by a different mechanism.

We note that participants’ baseline performance on two points discrimination task was very different across experiments. Several factors could account for this difference. One potential issue is that the tools and procedures used to test two-point discrimination (standard and digital caliper) differed slightly across (but not within) experiments. A second factor is that there are individual differences in tactile perception. However, we note that the effects observed remained when controlling for baseline accuracy, suggesting that these factors could not explain our results. Furthermore, the present study tested the effect on the increase in the perceived size induced in the somatosensory modality focusing of a specific body part, the lips. Similar results could be obtained testing other body parts, and future studies should test this hypothesis directly.

To conclude, although the mechanisms underlying the improvement in two point discrimination after AC remain speculative, a major contribution of the present work is to show that the body representation is dynamic and may be manipulated in minutes by altering sensory input. Our results represent the first evidence that the magnification of a body part derived from somatosensory inputs also has a beneficial effect on tactile perception.

## Materials and Methods

### Methods Experiment 1

Twenty adults (mean age = 23.4, *SD* = 4.9; 12 females) were tested with a two-point discrimination task (2PD; Haggard et al., 2003) in which they were asked to determine if the lower lip was touched with one or two points. During the task, participants closed their eyes. The task was performed three times in one session: (1) prior to application of AC (PRE condition), (2) ∼2 minutes after application of AC on both upper and lower lip (Lanacane, benzethonium chloride 0.2% and 20% benzocaine) (AC condition), and (3) ∼2 minutes after the AC was removed (POST condition).

In each administration of the task, 90 trials were presented. On 45 trials, a single touch was presented; on the other 45 trials, subjects were touched with two points spaced at a distance of 1, 2 or 3 mm (15 trials each). In each of the three conditions, these stimuli were applied manually at the center of the lower lip using a two-point discriminator (Baseline 12-140 Aesthesiometer, Fabrication). Subjects were instructed to indicate if their lips were touched by one or two points by depressing the appropriate key on a button box. The task was programed using E-Prime 3.0 software (Psychology Software Tools, Pittsburgh, PA) to randomize the order of the stimuli and of the response keys. Ninety trials were completed in approximately 15 minutes.

Subjects were also asked to judge the effect of the anesthetic cream on the size of their lips. Before each condition, participants were shown 8 drawings of a face (25 x 30 mm) on which the lips were depicted in sizes ranging from 5 x 2 mm to a maximum of 7 x 5.5 mm, with a vertical increase of about 0.5 mm, arranged in a 2×4 array from small to large size; subjects were asked to point to the face that depicted the size of their lips at that moment in time (see Figure 7 left panel).

**Figure 7:**
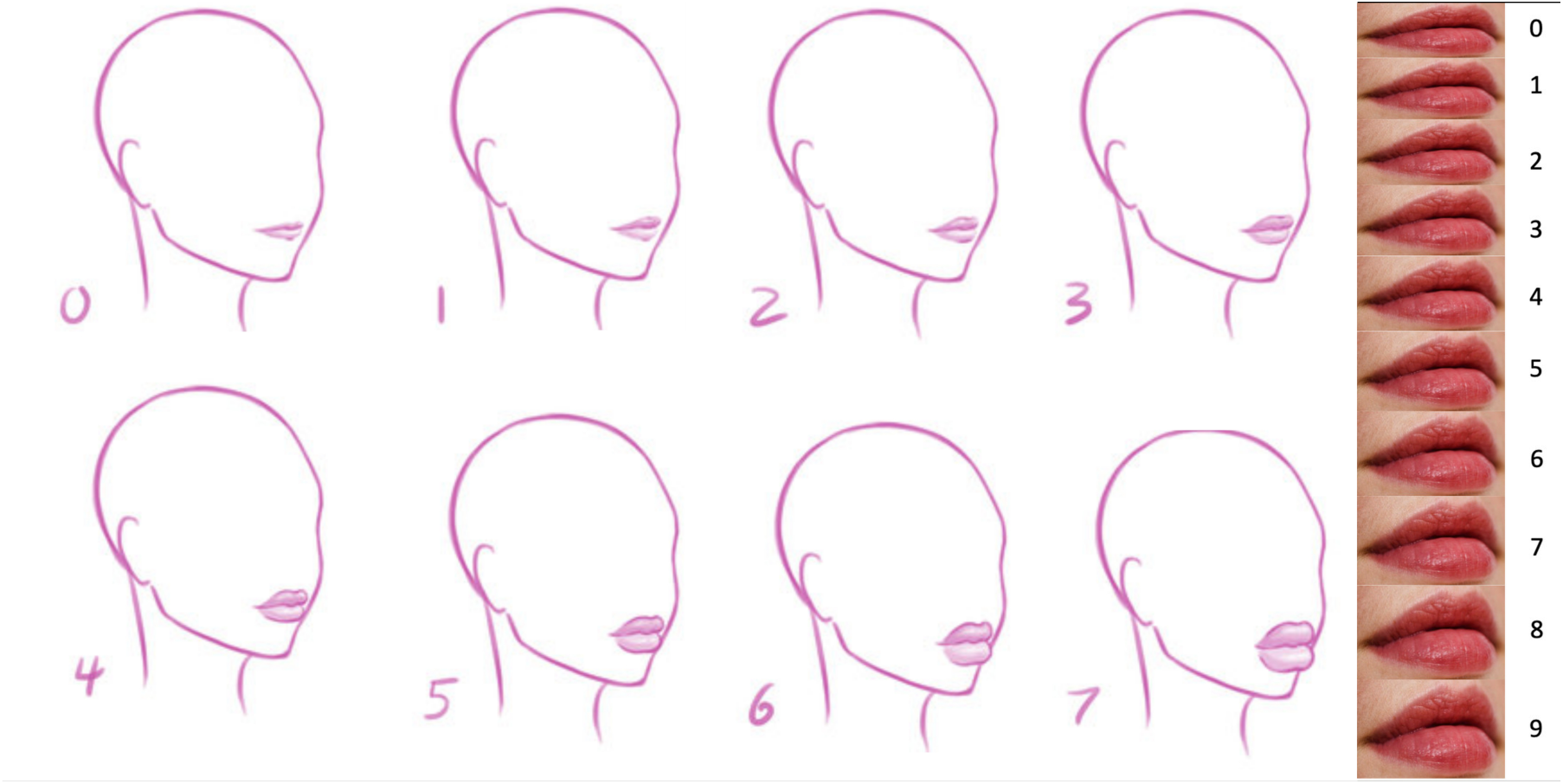
Representation of the stimuli used for the perceived size ratings.

### Methods Experiment 2

Forty individuals (mean age = 21.8, *SD* = 3.4; 29 females) underwent two testing sessions: in one session the same anesthetic cream as in Experiment 1 was used, while no cream was applied in the other session. Other differences with Experiment 1 included the following: (i) two-point discrimination was measured with an electric aesthesiometer (AOS ABSOLUTE Digimatic Caliper; resolution 0.01mm); (ii) tactile detection threshold was assessed with Semmes-Weinstein monofilament (Stoelting) and subjective rating of numbness in the PRE, EXPERIMENT and POST condition in both sessions, using a visual analogue scale from 0 (no numbness) to 10 (high numbness). Detection threshold was established by identifying the thinnest filament that the subject correctly detected on 2 of 3 trials. The procedure was initiated with a 5.8 nN filament and if they responded correctly on 2 of 3 trials, the next thinnest filament was presented.

### Methods Experiment 3

Fourteen individuals (*M*_age_ = 22.07, *SD* = 2.7; 9 females) underwent two testing sessions: in one session AC was applied on participants’ lips in one session, while in the other session MC (Eucerin^TM^) was applied. Participants were not told which cream was applied in each session and the order of sessions was counterbalanced between participants. The experimenter applying the tactile stimuli was blind to the EXPERIMENT condition. An electric aesthesiometer (AOS ABSOLUTE Digimatic Caliper; jaws resolution 0.01mm) was used to test two-point discrimination. Tactile detection threshold was assessed with Semmes-Weinstein monofilament (Stoelting), and the numbness was rated using a visual analogue scale from 0 (no numbness) to 10 (high numbness) in the PRE, EXPERIMENT and POST conditions.

### Data analyses

To test if the lip numbness ratings and tactile detection scores changed across time points (PRE, EXPERIMENT, POST) and conditions (no AC-AC in Experiment 2 and MC-AC in Experiment 3), we ran separate repeated measures ANOVAs. Post hoc t-tests are carried comparing difference (EXPERIMENT -PRE) in our dependent variable for between conditions.

In all Experiments, accuracy data from the two-point discrimination task was analyzed using logistic mixed effect model (LMM) in R (3.3.0) with subjects included as random intercepts. Factors under investigation and interactions between them were inserted into the model sequentially and ANOVAs were used to test the difference between models with or without the inclusion of each factor. Only factors contributing significantly to the model fit were included in the final model. Omnibus of the model were further estimated using the pamer.fnc function in R. We opted for this type of analysis because it accounts for the variability in the performance across individuals (Kliegl, Wei, Dambacher, Yan, & Zhou, 2011) and reduces the possibility of Type I error (Judd, Westfall, & Kenny, 2012). As our hypotheses and comparisons were specified *a prior*i specified, analyses were not adjusted for multiple comparisons.

Participants who performed 2 SD below the mean of the overall sample in the PRE condition were excluded from the analysis (one participant in Experiment 1 and two in Experiment 2)-

The Institutional Review Board of the University of Pennsylvania approved the present study and all participants signed an informed consent. The dataset of the current study is available from the corresponding author upon request. This study was carried out in accordance with the Declaration of Helsinki.

## Acknowledgements

We would like to thank Nicole White for her suggestions regarding the statistical analysis. We also would like to thank Marlee Coyle, Samuel Cason, Shayna Goldstein, and Eirini Zoupou for their help in testing participants. The study was supported by an R21 grant from NINDS awarded to H.B. Coslett. The authors declared that they had no financial interests or conflicts of interest with respect to their authorship or the publication of this article.

## Author contributions statement

E.A. designed the experiment, tested participants, analyzed the data and wrote the manuscript and H.B.C designed the experiments and revised the manuscript.

