## Supplementary material for "Apparent increase in lip size is linked to tactile discrimination improvement": SI

**Supplementary Information**

**Results Experiment 1**

The percentage of yes responses was similar in PRE and POST conditions, except for 2 ($z$=4.7, $p$<0.001) and 3mm ($z$=2.4, $p$=0.01).

For the POST condition, the difference in the lips size did not contributed to the model fit for whole face (logLik= -706, $\chi^2$ (4)=2.2, $p$=0.68) measure, but for the lips only (logLik= -7062 $\chi^2$ (4)=10.7, $p$=0.02). However, when further exploring this interaction, the effect of the size did not reached significance for any of the distances.

The LLM analysis exploring the effect of the perceived lips size was also carried out on the difference in the lips size based on the lips only rating and showed similar results as the whole face rating. The effect of the difference in lips size contributed significantly to the model fit for the EXPERIMENTAL condition (logLik= -684, $\chi^2$ (4)=16.5, $p$=0.002) (see SI Figure 1) and yes responses increased as function of the perceived changes in lips size at 1,2, and 3 mm (in all comparisons $z$>2.8, $p$<0.01). For the POST too the difference in lips size contributed significantly to the model fit for yes responses (logLik= -702, $\chi^2$ (4)=10, $p$=0.03), but post hoc test did not show any significant effect of the change in the lips size for any of the distances.

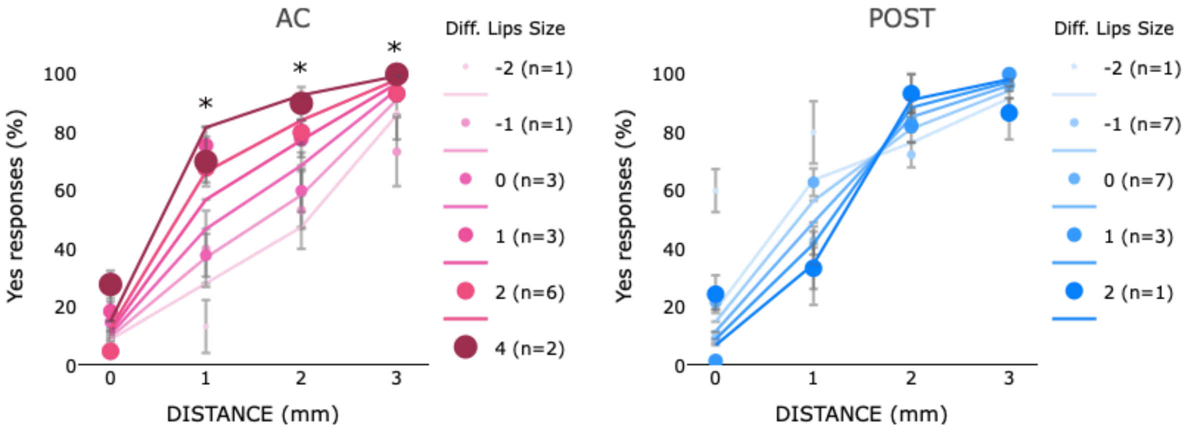


**SI Figure 1:** Mean (markers) and SE of the means (error bars) of yes responses in two points discrimination in AC (left panel) and MC (right panel) conditions as function of the distance (0,1,2,3 mm) between 2 points and the change in the perceived lips size (lips only rating) in Experiment 1. The different size/coloring of the markers described the performance of individuals based on change in the perceived lips size in that condition with respect to the PRE. The legend describes the amount of perceived change in the lip size and the number of individuals who perceived that change. Negative numbers indicate a perceived decrease and positive numbers indicate an increase in the perceived lip size compared to PRE. In every plot, colored lines represent the estimated effects in the LMM analyses.

**Table2.** Omnibus of logistic mixed effect model analysis.

| | Analysis 1: Effect of AC | Analysis 2: Effect of the perceived lips (lips only) | Analysis 2: Effect of the perceived lips (whole face) |
|---|---|---|---|
| **EXPERIMENT 1** | **Time:** *F=0.41, p=0.6, semi-partial R2= 0.29*<br>**Distance:** *F=449, p<0.001, semi-partial R2= 0.99*<br>***Time x Distance:*** *F=8.07, p<0.001, semi-partial R2=0.89* | **EXPERIMENTAL:**<br>**Lips size:** *F=4.9, p=0.02, semi-partial R2= 0.83*<br>**Distance:** *F=482, p<0.001, semi-partial R2= 0.99*<br>**Lips size *x Distance:*** *F=12.1, p=0.007, semi-partial R2=0.80.*<br>*Increase in yes as function of size for 1,2,3, mm (z>2.3, p<0.05)*<br><br>**POST:**<br>**Lips size:** *F=0.39, p=0.53, semi-partial R2= 0.31*<br>**Distance:** *F=158, p<0.001, semi-partial R2= 0.99*<br>**Lips size *x Distance:*** *F=3.2, p=0.02, semi-partial R2=0.76*<br>*The size did not reach significance for any distance* | **EXPERIMENTAL:**<br>**Lips size:** *F=9.6, p=0.001, semi-partial R2= 0.90*<br>**Distance:** *F=166, p<0.001, semi-partial R2= 0.99*<br>**Lips size *x Distance:*** *F=0.4, p=0.74, semi-partial R2=029*<br>*Increase in yes as function of size for 1 (z=3.1, p=0.001) 2, mm (z=2.7, p=0.02)*<br><br>**POST:**<br>**Lips size:** *F=0.16, p=0.16, semi-partial R2= 0.14*<br>**Distance:** *F=164, p<0.001, semi-partial R2= 0.99*<br>**Lips size *x Distance:*** *F=0.70, p=0.02, semi-partial R2=0.41* |
| **EXPERIMENT 2** | **Time:** *F=2.3, p=0.09, semi-partial R2= 0.70*<br>**Distance:** *F=1491, p<0.001, semi-partial R2= 0.99*<br>**Condition:** *F=6.9, p=0.008, semi-partial R2= 0.87*<br>**Time x Distance:** *F=0.90, p=0.48, semi-partial R2= 0.47*<br>**Time x Cream** *F=5.4, p=0.003, semi-partial R2= 0.84*<br>**Distance x Cream** *F=5.8, p<0.001, semi-partial R2= 0.85*<br>***Time x Distance x Cream:*** *F=2.0, p=0.054, semi-partial R2=0.67* | **AC:**<br>**Lips size:** *F=4.4, p=0.03, semi-partial R2= 0.81*<br>**Distance:** *F=256, p<0.001, semi-partial R2= 0.99*<br>**Lips size *x Distance:*** *F=1.0, p=0.36, semi-partial R2=0.51.*<br><br>**No AC**<br>**Lips size:** *F=0.42, p=0.51, semi-partial R2= 0.30*<br>**Distance:** *F=254, p<0.001, semi-partial R2= 0.99*<br>**Lips size *x Distance:*** *F=1, p=0.36, semi-partial R2=0.51.* | **AC:**<br>**Lips size:** *F=2.1, p=0.14, semi-partial R2= 0.68*<br>**Distance:** *F=256, p<0.001, semi-partial R2= 0.99*<br>**Lips size *x Distance:*** *F=0.34, p=0.79, semi-partial R2=0.26*<br><br>**No AC**<br>**Lips size:** *F=0.42, p=0.83, semi-partial R2= 0.04*<br>**Distance:** *F=254, p<0.001, semi-partial R2= 0.99*<br>**Lips size *x Distance:*** *F=0.79, p=0.49, semi-partial R2=0.44* |
| **EXPERIMENT 3** | **Time:** *F=4.1, p=0.01, semi-partial R2= 0.80*<br>**Distance:** *F=576, p<0.001, semi-partial R2= 0.99*<br>**Condition:** *F=2.8, p=0.09, semi-partial R2= 0.73* | **AC:**<br>**Lips size:** *F=0.14, p=0.70, semi-partial R2= 012*<br>**Distance:** *F=87, p<0.001, semi-partial R2= 0.99*<br>**Lips size *x Distance:*** *F=6, p<0.001, semi-partial R2=0.85.*<br><br>**MC** | **AC:**<br>**Lips size:** *F=0.78, p=0.37, semi-partial R2= 044*<br>**Distance:** *F=87, p<0.001, semi-partial R2= 0.98*<br>**Lips size *x Distance:*** *F=5.5, p<0.001, semi-partial R2=0.44.*<br><br>**MC** |

| | | | |
|---|---|---|---|
| | **Time x Distance:** *F=3.8, p=0.008, semi-partial R2= 0.79*<br>**Time x Cream** *F=0.53, p=0.58, semi-partial R2= 0.34*<br>**Distance x Cream** *F=3.8, p=0.008, semi-partial R2= 0.79*<br>***Time x Distance x Cream:*** *F=3.2, p=0.003, semi-partial R2=0.76* | **Lips size:** *F=0.02, p=0.87, semi-partial R2= 0.02*<br>**Distance:** *F=77.6, p<0.001, semi-partial R2= 0.98*<br>**Lips size *x Distance:*** *F=0.1, p=0.93, semi-partial R2=0.11.* | **Lips size:** *F=0.60, p=0.43, semi-partial R2= 0.02*<br>**Distance:** *F=76.4 p<0.001, semi-partial R2= 0.98*<br>**Lips size *x Distance:*** *F=0.58, p=0.62, semi-partial R2=0.11.* |

**Results Experiment 2**

The LLM analysis with condition (AC or no AC), time point (PRE, EXPERIMENTAL, and POST) and distance (0, 1, 2 and 3 mm) showed an effect of the condition in the PRE. Indeed, performance in the AC was better than the no AC at 2mm (z=2.1, p=0.03) and 3 mm (z=2.9, p=0.03); while no significant differences between these two conditions were observed in the POST. To test if the performance at baseline could account for the observed difference in the EXPERIMENTAL condition, we carried out an additional LMM analysis on this condition inserting the *yes* responses at baselines as a fixed factor. This factor did not improve the model fit with respect to the model with distance and condition as fixed factor (logLik= -3534, $\chi^2$ (8)=10.8, $p$=0.20) and its inclusion did not change our results, suggesting that the variability in the baseline performance across condition could not account for the difference observed in the EXPERIMENTAL condition. Furthermore, for the AC, performance the EXPERIMENTAL condition was better than the POST for 1 ($z$=4, p<0.001) and 2 (z=3.6, p<0.001) mm; no significant differences were observed between POST and EXP. For the No AC, performance was consistent across time points for all the distances.

The LLM analysis exploring the effect of the perceived lips size was also carried out on the difference in the lips size based in the whole face rating. The effect of the difference in lips size did not contribute significantly the model fit (logLik= -1734, $\chi^2$ (4)=3.1, $p$=0.53) (SI Figure 2). This was also true for the no AC condition (logLik= -1767, $\chi^2$ (4)=2.5, $p$=0.64).

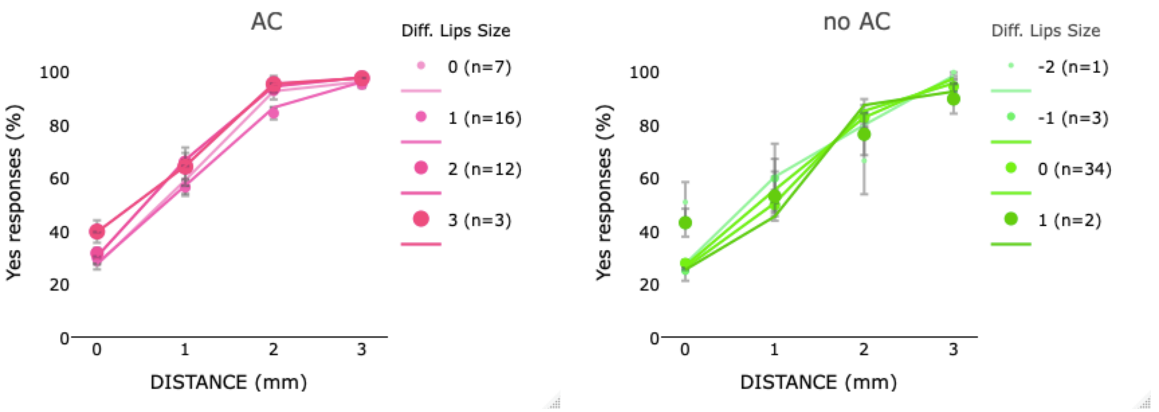


**SI Figure 2:** Mean (markers) and SE of the means (error bars) of yes responses in two points discrimination in AC (left panel) and MC (right panel) conditions as function of the distance (0,1,2,3 mm) between 2 points and the change in the perceived lips size (whole face rating) in Experiment 2. The different size/coloring of the markers described the performance of individuals based on change in the perceived lips size in that condition with respect to the

PRE. The legend describes the amount of perceived change in the lip size and the number of individuals who perceived that change. Negative numbers indicate a perceived decrease and positive numbers indicate an increase in the perceived lip size compared to PRE. In every plot, colored lines represent the estimated effects in the LMM analyses.

**Results Experiment 3**

The LLM analysis with condition (AC or no AC), time point (PRE, EXPERIMENTAL, and POST) and distance (0, 1, 2 and 3 mm) showed an effect of the condition in the PRE. The percentage of *yes* responses was higher in MC than AC at 1mm ($z$=2, $p$=0.04) and in the AC than MC at 3mm (z=2.3, $p$=0.01) (see SI Figure 3). As in Experiment 2, we examined if the performance in the PRE conditions could account for the results observed in the EXPERIMENTAL and POST condition; to address this point, we ran a LMM on accuracy in these two conditions and tested the effect of baseline accuracy. This factor did not contribute significantly to the model fit and its inclusion did not change our results. No significant difference between AC and MC was observed in the POST.

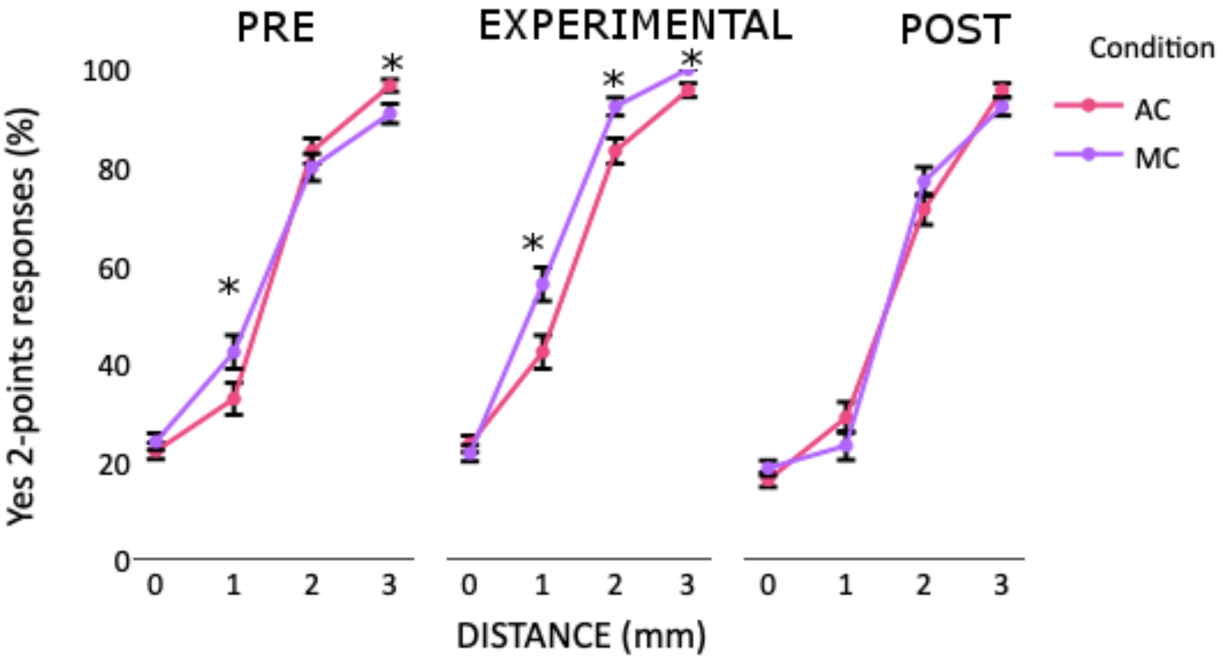


**SI Figure 3:** Mean (markers) and SE of the means (error bars) of yes responses in two points discrimination in PRE, EXPERIMENT and POST conditions for AC and no AC conditions at each distance (0,1,2,3 mm) in Experiment 2.

The LLM analysis exploring the effect of the perceived lips size was also carried out on the difference in the lips size based on the lips only rating. The effect of the difference in lips size contributed to the model fit of yes responses for AC (logLik= -567, $\chi^2$ (4)=19, $p$<0.001), but not for MC (logLik= -517, $\chi^2$ (4)=0.4, $p$=0.9). *Yes* responses increased as a function of perceived increase in lip size only for AC, at 3 mm ($z$=2.4, $p$=0.01).

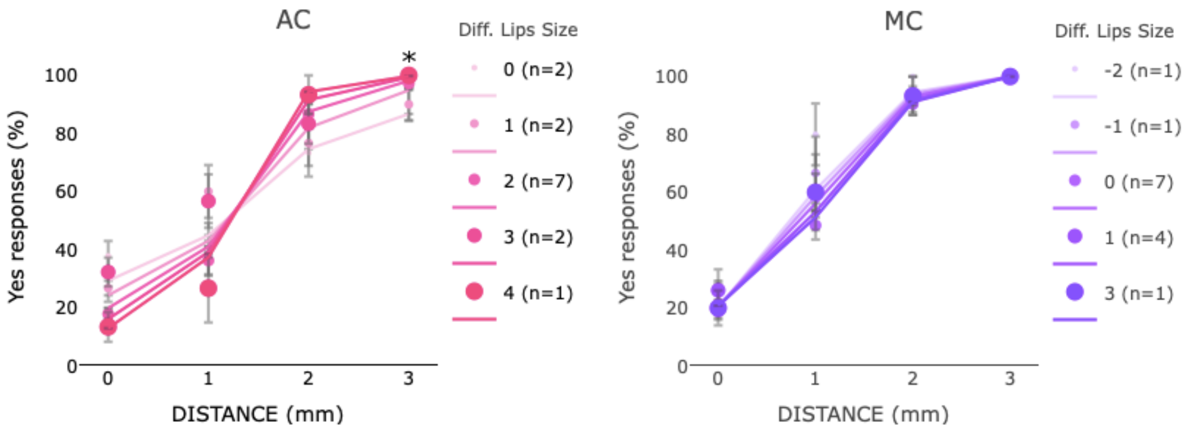


**SI Figure 2:** Mean (markers) and SE of the means (error bars) of yes responses in two points discrimination in AC (left panel) and MC (right panel) conditions as function of the distance (0,1,2,3 mm) between 2 points and the change in the perceived lips size (lips only face rating) in Experiment 3. The different size/coloring of the markers described the performance of individuals based on change in the perceived lips size in that condition with respect to the PRE. The legend describes the amount of perceived change in the lip size and the number of individuals who perceived that change. Negative numbers indicate a perceived decrease and positive numbers indicate an increase in the perceived lip size compared to PRE. In every plot, colored lines represent the estimated effects in the LMM analyses.
